# Development of predictive QSAR models on the phosphopeptide binding affinity against 14-3-3 isoforms

**DOI:** 10.1101/2020.07.24.217752

**Authors:** Ying Fan, Xiaojun Wang, Chao Wang

**Affiliations:** Shenzhen Institute of Information Technology, Shenzhen, 518172, China; HKBU Institute for Research and Continuing Education, Shenzhen 518057, China

**Keywords:** 14-3-3s, bioinformatics, QSAR, peptide microarray

## Abstract

14-3-3s present in multiple isoforms in human cells and mediate signal transduction by binding to phosphoserine-containing proteins. Recent findings have demonstrated that 14-3-3s act as a key factor in promoting chemoresistance of cancer. Here, we aimed to develop the predictive models that can determine the binding affinity of phosphopeptide fragments against 14-3-3s. It is found that the hydrophobic property of residues in phosphopeptides has a significant contribution to the binding affinity for most of 14-3-3 isoforms. The conserved patterns of 14-3-3 biding motifs were verified by our predictions. A group of peptide sequences were predicted with high binding affinity and high sequence conservation, which had agreement with 14-3-3s ligands. Overall, our results demonstrate that how the residues are likely to function in 14-3-3s interaction and the computational methods we introduced may contribute to the further research.

## 1 Introduction

14-3-3 proteins constitute a family of highly conserved acidic proteins with the size of around 30 kDa [2, 53, 1, 4]. These proteins express in high abundance in a wide range of eukaryotic cells and exist in many tissues [16]. It is found that 14-3-3s present in multiple isoforms in human cells and function as either homo- or hetero-dimer formation [16, 38]. The dimer presents a conserved groove that served as the phosphopeptide binding pocket and positively charged residues (e.g. K49, R56, and R127) in the pocket interact with the phosphorylated binding partners/ligands. 14-3-3s binding to specific phosphoserine-containing motifs could result in the assembly of important signaling complexes [39]. The interactions of 14-3-3s with signaling proteins are critical for the activation of signal transduction processed [11]. All of these processes involve multiple isoforms of 14-3-3. Through interaction with their binding partners, 14-3-3s involve in the control of diverse cellular processes, including metabolism, transcription, cell cycle, differentiation, migration and apoptosis [50, 25]. Recent findings have demonstrated that 14-3-3s as a key factor in promoting chemoresistance of cancer cell [40, 46, 51, 19]. 14-3-3s mediated tumorigenesis and chemoresistance by gene overexpression has been observed by the animal studies [42]. Also, high expression level of 14-3-3s was found to be associated with disease recurrence and poor survival in the cancer patients received chemotherapy [32, 59, 22]. These evidences suggest that 14-3-3s involve in chemoresistance in multiple human cancers as key regulators. This significant association of 14-3-3s overexpression with poor patient outcomes makes this protein an attractive candidate targets for effective cancer management. Hence, 14-3-3s can be a novel biomarker for the diagnosis and a potential therapeutic target in cancer therapy.

The human 14-3-3 protein family consists of seven distinct but related isoforms, which are named *β, γ, ϵ, η, σ, τ, ζ*. The seven isoforms are highly conserved in sequence and structure, each expressed by a particular gene [53]. Hence, there is functional overlap between 14-3-3s, and the ability of one isoform to compensate for the loss of another has been demonstrated in some cases [55]. A comparison of crystal structures from different human 14-3-3s shows striking structural similarities [18]. However, the distinct differences existed for each 14-3-3 isoform existed in the preference of the dimerization and binding specificity [18]. Isoform *γ* have both two kind of dimeric formations, but mainly with *ϵ* for heterodimers [55]. Only heterodimer but not homodimer formation is found in the isoform *ϵ* [52, 55]. Both homo- and hetero-dimeric formations have been found in 14-3-3s. One exception is that isoforms *σ* prefer to form homodimers [55, 3, 18]. These preferences derive from the slightly structural and sequence variances. Also, the binding partners of 14-3-3s show a distinct preference for a particular isoform. The studies to determine 14-3-3s binding sites reveal that 14-3-3s bind to phosphoserine/threonine (pS/T) in a sequence-specific manner [60, 36]. There are three modes of motifs found in 14-3-3s binding sites: mode-1 (RSXpSXP), mode-2 (PXF/YXpSXP) and mode-3 (SWpTX) at C-terminal [28]. While 14-3-3s are known as the family of phosphor-binding proteins, several exceptions related to out of motif or even without phosphor-specific manner were also identified [28]. The motif of phosphoserine-dependent interaction is important for high affinity 14-3-3s binding. This indicated that both phosphoserine and its flanking residues take part into the binding of 14-3-3s with its target ligand.

The binding affinity of 14-3-3s with its ligand can be obtained by high throughput combinatorial peptide microarray method [31]. Upon this, the 14-3-3s binding specificity to a wide variety of phosphopeptides can be determined in an experiment. But there is a lot of limitations exits still. For example, for a tripeptide, the number of possible sequence arrangements are 20^3^. Owing to the extremely large amount of possible arrangements, predicting the binding affinity of 14-3-3s with its ligand may be a challenging research topic in life science. Therefore, it is vitally important to develop a computational method for efficiently predicting peptides. At sequence level, the quantitative structure-activity relationship (QSAR) have been broadly used to infer the biological function and to predict high bioactivity peptide [10]. One of QSAR based approach is amino acid-based peptide prediction, in which the QSAR is used for the protein and peptide activity prediction [10]. For a given 14-3-3 protein, the property of binding affinity can be seen as a kind of bioactivity of the peptide sequence. In QSAR studies, linear regression has been commonly used to link the biological activities as a response variable to the molecular descriptors as predictor variables for data analysis [65]. Variable selection using penalized methods plays a vital role in statistical modeling with high dimensional QSAR data [49]. It aims to select only a subset of important descriptors from a large number of molecular descriptors, and thereby to improve the performance of QSAR models in terms of obtaining higher prediction accuracy of the model and easy interpretation. Penalized regression methods provide an estimate QSAR model that has lower prediction errors than multiple linear regression [44].

The details of the interaction between 14-3-3s and its ligands are not fully understood still, including the isoforms specificity, sequence patterns of the ligands, and the structural features of the interaction. Considering there are a great number of proteins that found to bind 14-3-3s *in vitro* and *in vivo*, it is meaningful to determine 14-3-3s binding interactions at large scale. Here, the computational predictive models to identify the binding affinity of the phosphopeptide sequences in seven 14-3-3 isoforms were reported. The raw affinity data was retrieved for literatures and processed before analysis. The divide physicochemical property scores (DPPS) were applied to generate variables matrix. The QSAR models were developed by linear regression method for phosphopeptide sequence prediction. The prediction of 14-3-3s binding affinity based on the data integrated with QSAR modelling. Based on this, target-specificity of seven mammalian 14-3-3 proteins were determined and the putative 14-3-3 motifs were identified.

## 2 Results

The purpose of this study is to develop the predictive QSAR models that can determine the binding affinity of phosphopeptide fragments against seven mammalian 14-3-3 isoforms. It helps us to decode the ligand interaction at residue level. The dataset contained the raw affinity values for 14-3-3s with phosphopeptide fragments was transformed and processed. The variables for describing the binding affinity of phosphopeptide fragments were extracted by DPPS. Penalized regression method was utilized to perform QSAR study and the models for seven isoforms were created by elastic net. Based on the established QSAR models, the contributions of various residues were analyzed. Finally, we evaluated the relative binding affinity of all possible sequence arrangements from the N- and C-terminal.

### 2.1 Dataset

Although the overall binding profiles of seven 14-3-3 isoforms with the phosphopeptide fragments library were quite resemble, there were distinctive features that differed from one isoform to another found, which shown a non-normal distribution and resulting in ineffective discrimination of different peptide sequences. Before modeling, all the data are pre-processed, and quality checked. First of all, the relative fluorescent values were log transformed and the peptide sequences with relative low values were removed from the library. Even the raw data is imbalance distributed, the transformed affinity data are normally distributed and but not consistent for each sub-library (Fig.1 A). Among of that, the affinity of isoform *ζ* is higher than the rest of six isoforms. After pre-processed, the mean values of affnity range from 5 to 6 for the seven isoforms. The pre-processed data are used for the rest of the analysis. From the analysis of high binding affinity fragments in this library, it could be found that the frequency of occurrence of a specific amino acid was different at each position. Similar finding can also be found in other reports [60].

**Figure 1:**
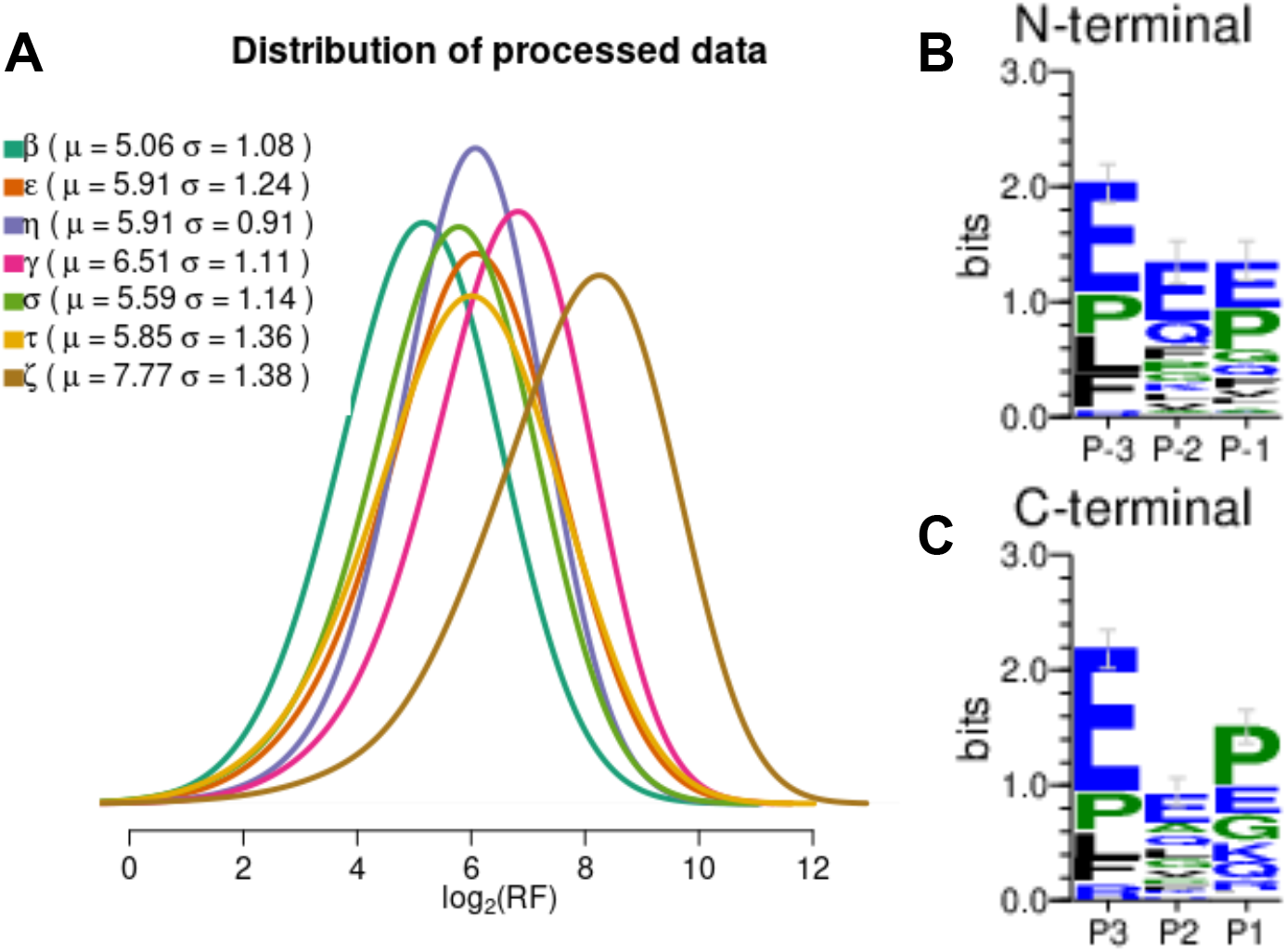
(A) The distribution of the binding affinity data after processing. The mean (*μ*) and standard deviation (*σ*) values for the seven isoforms are listed in the legend of the plot. (B) and (C) The conserved peptide sequences in N- and C-sublibrary with low binding affinity against 14-3-3s.

Totally 482 of log-transformed data that shown low binding affinity (less than zero) are discarded (210 in N-terminal sublibrary and 272 in C-terminal sublibrary). The low binding affinity value indicates the corresponding peptide sequence has no contribution in protein interaction, which could cause negative effects on model development. A small portion of sequences with low affinity was found (less than 7%) for each isoform except the isoforms *γ* (15.3%) and *σ*(15.0%). Most of the peptide sequences with low binding affinity exist only in one or two isoform libraries. Only three peptide sequences commonly exist in six isoform libraries, which is “EEP*p*SXXX” in N-terminal library, “XXX*p*SPEE” and “XXXpSPLE” in C-terminal library. And the multiple alignments shown that the peptide sequences in low affinity data are conserved. The 384 out of 482 peptide sequence have at least one Glu residue at any positions (Fig.1 B and C).

### 2.2 QSAR modeling on the relative affinity data

All of the 30 variables derived from DPPS descriptors were used to characterize each phosphopeptide fragments. For each sublibrary, 80% fragments with the relative binding affinity data from the experiments are treated as training data. The details on the identification of the relative binding affinity for all isoform sublibraries are shown (Fig.2). It clearly shows the robustness of the QSAR model developed in this study. For the 14-3-3 isoform *ζ*, the modelling results has overall *R*^2^ and *RMSE* values of 0.7897926 and 0.619047 in the N-terminal sublibrary, 0.7444364 and 0.7037071 in the C-terminal sublibrary. The *R*^2^ and *RMSE* values vary between seven isoforms. This might due to the fact that the relative binding affinity values distribute differently in the peptide library.

**Figure 2:**
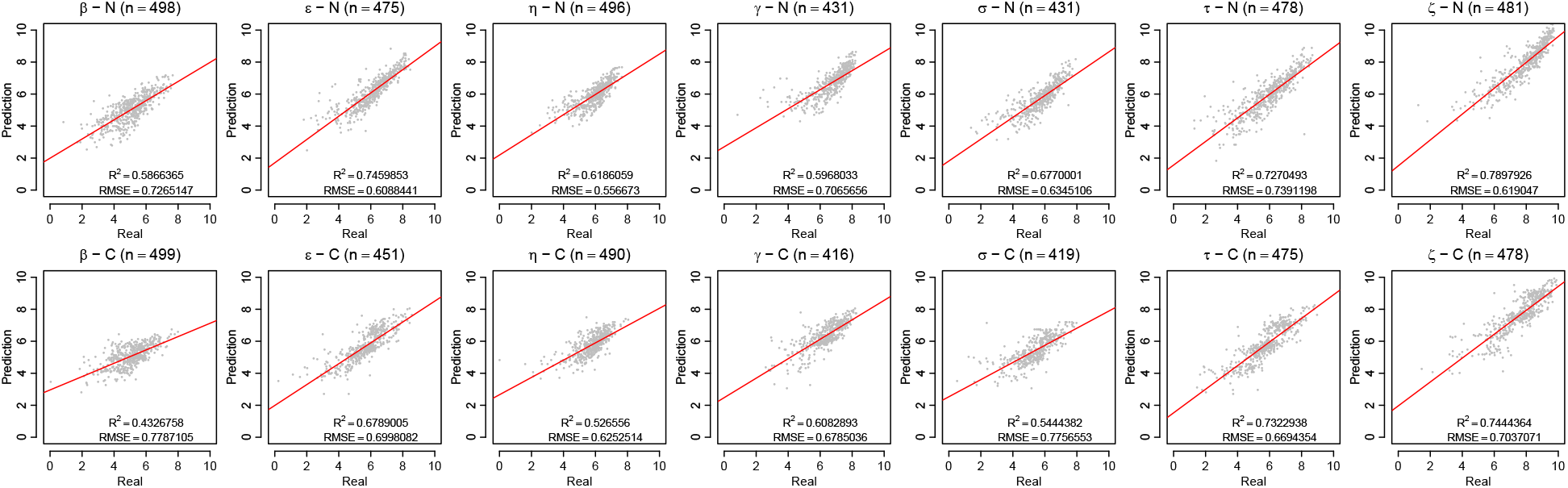
Scatter plot of the relative binding affinity values (real) versus prediction by the QSAR model (prediction) of 14-3-3 seven isoforms with the phosphopeptide fragments library. The title of the plot represents the isoform type and the sublibrary type of the data used in the analysis. The overall *R*^2^ and *RMSE* values are shown in the inset of the plots.

The coefficients of the QSAR models were used to explore the contributions of amino acid residues. The overall contribution of every physicochemical parameter at each position based on the normalized coefficients of the QSAR model was calculated and plotted to explore the preferred residue for 14-3-3 interaction (Fig.3). Relative to phosphoserine, the contribution of position −2 and −1 is associated with electronic property of residues, in addition to hydrophobicity. At position +1, the contribution also come from the partial of electronic property and hydrophobicity. Meanwhile, it can be found that the contribution is not common between seven isoforms. For example, electronic property at position −2 was only advantageous for the isoform *σ, τ*, and *ζ* to form high binding affinity. Arg and Lys were preferred at the position −1 relative to *pS*. And nonpolar residues are preferred at the position −1 and +1.

**Figure 3:**
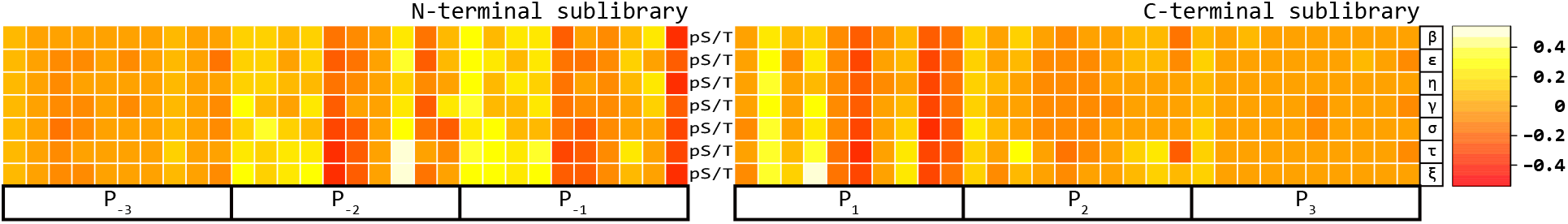
Heatmap of regression coefficients with N- and C-terminal sublibrary. For each amino acid residue position P, there are 10 coefficients corresponding to four types of physiochemical properties, which are electronic property (1-4), steric property (5-6), hydrophobicity (7-8) and hydrogen bond (9-10). The color scale of the heatmap is illustrated at right side.

The performance of the QSAR models for each isoform are evaluated by two ways. First, the consensus sequences of amino acids from the high binding affinity phosphopeptide fragments are identified. There is significant preference for Arg in position −3 relative to pS. The same selectivity of amino acid can be found in position −1 in addition to Lys. Phe and Ala is evidently selected in the position −2. For the C-terminal side, the slightly preference of amino acids are also determined. The selection for Ala in the position +1 and Phe in the position +2 is evident. These preferences of amino acids are highly consistent with the experiment data here we used and previous published work [60, 31]. Second, the distance and similarity of affinity among the seven isoforms are evaluated and compared with the experiment data. The distance of predicted binding affinity between the isoforms are changed slightly, which indicated that the accurate predictions are obtained by the models we developed.

### 2.3 Prediction of the relative binding affinity of peptides with 14-3-3s

The QSAR models developed from N- and C-terminal sublibrary against all seven 14-3-3 isoforms were used to predict the binding affinity of all possible tripeptide sequences (20^3^ = 8000) to 14-3-3s. The generated data was standardized by isoforms so that the mean was zero and the variance was one (Supplementary data). The standardized data was treated as the predicted results for the following analysis. The ranges of standardized data are −4.27 to 3.09 in N-terminal sublibrary and −4.46 to 3.67 in C-terminal sublibrary. Moreover, the mammalian proteins that have direct interactions with 14-3-3s from previous reports were collected. The relevant interaction sites with phosphopeptide were extracted and compared with the corresponding prediction results in this study (Tab.1). Disintegrin and metalloproteinase domain-containing protein 22, also know ADAM22, is a member of membrane protein in human [20]. The interaction of ADAM22 with 14-3-3s isoforms have confirmed by yeast two-hybrid screen [66]. There are two sites found to be conserved with consensus motif mode 1 in the peptide sequence. But both of these two sites are lack of Pro at position +2 relative to phosphoserine residue. This is consistent with the reports that an alanine substitution at +2 in synthetic peptide binding studies shows only a moderate effect on 14-3-3s binding [62]. On the basis of prediction in our results, it can be found that the imbalance binding affinity exist between N- and C-terminal.

**Table 1:**
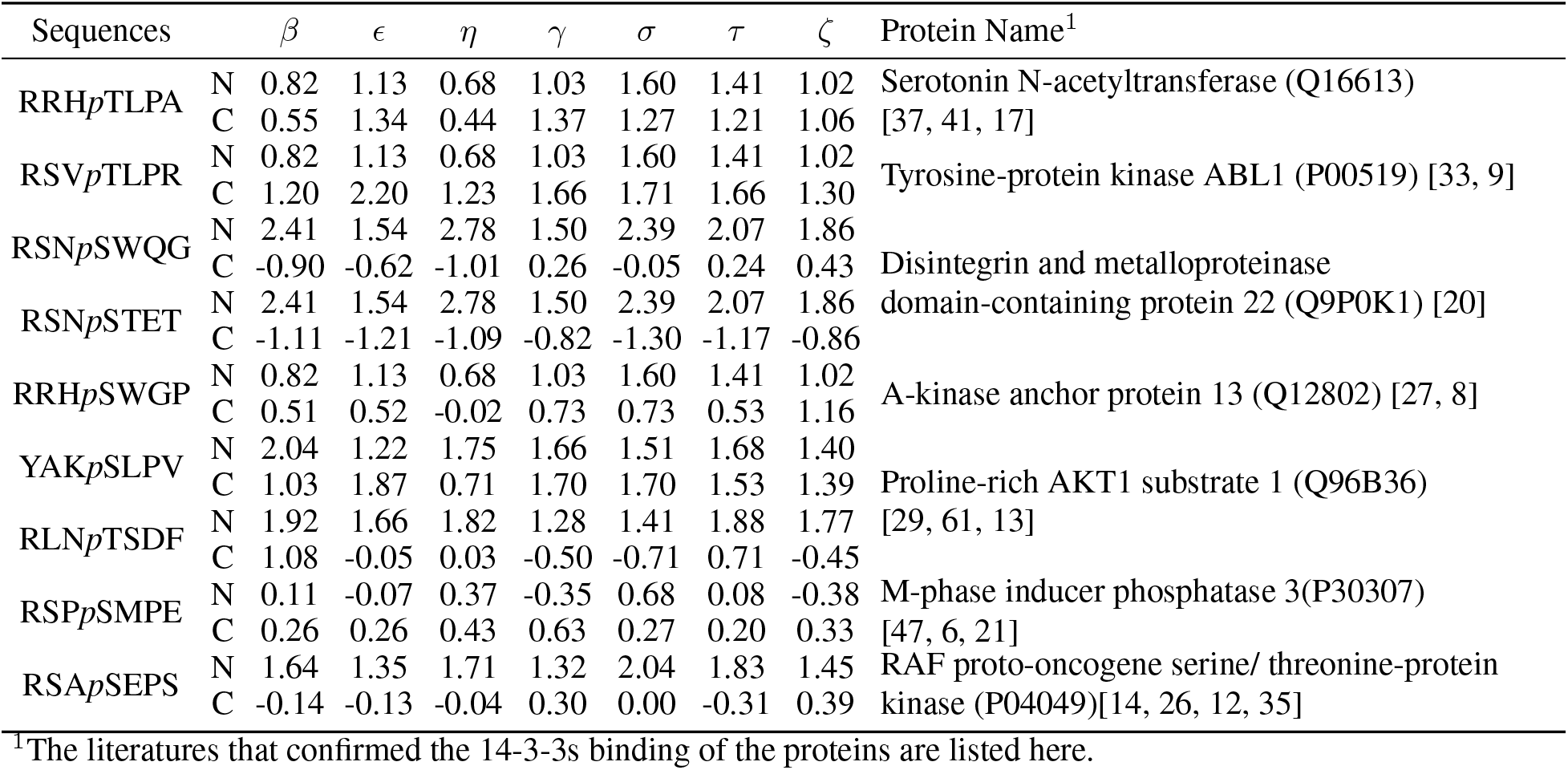
The predicted binding affinity of peptide sequences in the proteins that have direct interactions against 14-3-3s from literatures.

The QSAR models developed from N- and C-terminal sublibrary against all seven 14-3-3 isoforms were used to predict the binding affinity of all possible tripeptide sequences (20^3^ = 8000) to 14-3-3s. The generated data was standardized by isoforms so that the mean was zero and the variance was one (Supplementary data). The standardized data was treated as the predicted results for the following analysis. The ranges of standardized data are −4.27 to 3.09 in N-terminal sublibrary and −4.46 to 3.67 in C-terminal sublibrary. Moreover, the mammalian proteins that have direct interactions with 14-3-3s from previous reports were collected. The relevant interaction sites with phosphopeptide were extracted and compared with the corresponding prediction results in this study (Tab.1). Disintegrin and metalloproteinase domain-containing protein 22, also know ADAM22, is a member of membrane protein in human [20]. The interaction of ADAM22 with 14-3-3s isoforms have confirmed by yeast two-hybrid screen [66]. There are two sites found to be conserved with consensus motif mode 1 in the peptide sequence. But both of these two sites are lack of Pro at position +2 relative to phosphoserine residue. This is consistent with the reports that an alanine substitution at +2 in synthetic peptide binding studies shows only a moderate effect on 14-3-3s binding [62]. On the basis of prediction in our results, it can be found that the imbalance binding affinity exist between N- and C-terminal.

The predicted results confirm that high conserved binding affinity among seven 14-3-3 isoforms. The sequences of tripeptide RST (c-Raf-1, A-Raf), RDS (Cdc25a), RPS (Cdc25b), RAA (PKC-*ϵ*), RAK (PCTAIRE-2), RSH (mT), RHA (TH), RHS (TPH) and RSK (A20) at N-terminal sublibrary matched motif mode 1 all are shown high binding affinity (standardized affinity > 1.5), except RSP (Cdc25c). It was reported that 14-3-3 participate in the cell cycle regulation by interacting with Cdc25c [21]. The phosphopeptide sequence in the flanking regions of Cds25c residue 216 is RSP*p*SMPE, which partially match motif mode 1. Although the predicted binding affinity of RSP was low. It can be found that MPE at C-terminal present high binding affinity in our result, which suggested that the contribution of interaction of Cdc25c against 14-3-3s might be different with the other ligands. RIH (Cdc25a), RFQ (Cdc25b), CVR (PKC*γ*), PTR (IRS-1), SYT (CK-8), LYR (PICALM) at N-terminal sublibrary matched motif mode 2 are also shown high binding affinity in predicted results.

### 2.4 Cluster analysis of the relative binding affinity

The two-way hierarchical cluster analysis was utilized based on the normalized binding affinity data (Fig.4). The distinct clustering of the isoform-specific group was not found in the clustering result. However, the clustering identified a group of peptide sequences with high binding affinity in the N-terminal sublibrary. The number of tripeptides in the group are 253. The values of binding affinity in this group are significantly higher than the rest of data from all seven isoforms (*p* < 0.0005).

**Figure 4:**
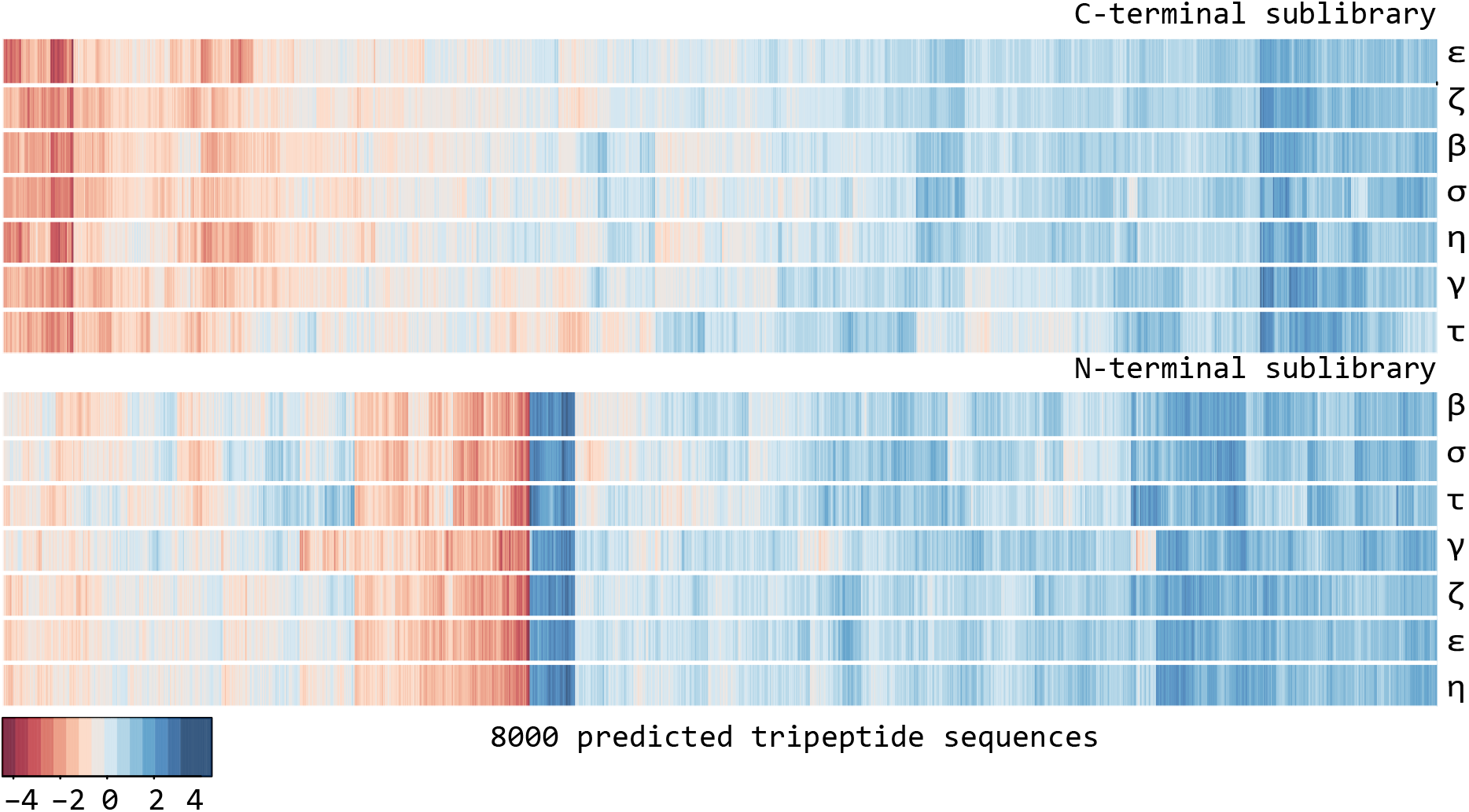
Hierarchical cluster analysis of the standardized binding affinity data precited by the QSAR models. The color scale is illustrated at the bottom. The group of sequences showed significant binding affinity is highlighted with black contours.

The pattern of sequences in this group are enriched in the binding motif mode 1. The tripeptide sequences that is commonly found from previous reports are clustered in this group, e.g. RDS, RST, RFQ, RAA, RAK, RSK, RPS and so on (Supplementary data). The position specific scoring matrix produced by the peptide sequences indicated that strong selection for peptides containing Arg or Lys in the position −3. This finding was consistent with the patterns found in the experiment dataset[31]. And non-polar amino acid at the position −2, e.g. Ala, Pro, Cys, Trp, Tyr, Phe, Ser. Ser, Phe or Tyr in the *p*S −2 position have been reported in the degenerate peptide library [60]. And in the position −1, Arg, Lys, Asn are the most common residues, which have been illustrated before[28].

## 3 Discussion

For the development of QSAR models, it is important to screen the amino acid descriptors. To date, there are diverse types of amino acid descriptors are available from published literatures or database, e.g. and so on. For improving the performance of modeling, several types of descriptors commonly used in the QSAR modeling were collected and used in this study to obtain a suitable descriptor. There descriptors were z-scales [24], VHSE[34], HESH [45], G8 [54] and DPPS[56]. It is commonly known that every type of descriptor has their own advantages and limitations. The z-scale is widely applied in and proven to be useful for the prediction of peptide activity [23]. But the limitations in the modeling of tripeptide sequences were also reported [57]. Although VHSE and z-scale were constructed from different kinds of physicochemical parameters, both of them have three types of variables (hydrophobicity, steric property and electronic property). Previous study found the different results generated using VHSE and z-scale [30]. And in the activity prediction of angiotensin-converting enzyme inhibitors, bradykinin-potentiating pentapeptides, bitter tasting dipeptides, the application of VHSE descriptors could acquire satisfactory results [54, 34]. This suggested the advantageous use of VHSE for describing the structural variability in the protein-protein interactions. More physicochemical parameters were included both by DPPS (*n* = 171) and HESH (*n* = 119) than VHSE (*n* = 50). The property of hydrogen bond was added into DPPS and HESH to describe the peptides. These two types of amino acid descriptors have been applied to build some good models with different regression methods [48,45,64]. Even DPPS and HESH were newly published descriptors, the quality of the models (*R*^2^ and *RMSE*) developed by HSEH and DPPS were not improved in our preliminary study. However, the performance of prediction with DPPS were better than the rest of descriptors in the preliminary results of this study. Thus, DPPS was selected as the amino acid descriptor here. At the same time, various regression methods (including support vector regression, random forest, partial least squares, multiple linear regression, lasso and ridge) were evaluated by the quality of the models in the preliminary studies. Even the *R*^2^ and *RMSE* of the models by these methods were totally different, it can be found that the models by elastic net method had the most powerful fitting capacity. Hence, elastic net was used here for developing the QSAR models.

The ten types of amino acids used in the peptide library only account for 50% of total, especially only five types at *P*±_3_. In this case, we could not get the full views on the 14-3-3s binding preference due to the limitations existed in the experiments. The QSAR modeling method provide an alternative way to extend the knowledge on 14-3-3s interactions. According to the coefficients from QSAR modeling, the important properties of residues were identified. The hydrophobic property of residues at N-terminal has a significant contribution to the binding affinity of phosphopeptides. The conserved patterns of phosphopeptide fragments in the ligands were verified in our predicted results. In addition, most of the proteins that reported to bind to 14-3-3s through experiment were also detected in our study. The binding affinity of the phosphopeptide sequences in these well know proteins were not determined yet by the combinatorial peptide library [31]. Our predictions provide the evidences that the QSAR modeling method can be applied to such type of research and insightful information could be obtained. Meanwhile, new peptide sequences with high affinity value were also determined, which have great potentials to be found at the next step of research. After being screened and validated by in vitro and in vivo studies, these peptides that was not found before are plausible to be developed as the inhibitors or agents that interfere with 14-3-3 related pathways and as novel therapeutic molecules for targeted cancer therapy with purpose of clinical research.

## 4 Materials and Methods

### 4.1 Dataset

The dataset was extracted from a fingerprint peptide library coupled with seven mammalian 14-3-3 isoforms [31]. The different source of isoforms was used in this peptide library. The 14-3-3s binding affinity data in the library was generated by the fragment-based combinatorial peptide microarray and consisted of 1000-member phosphopeptide fragments with seven 14-3-3 isoforms, respectively. The data was split into two sublibraries. There were 500 members of N-terminal *P*_−3_*P*_–2_*P*_−1_-*p*S-*X*_+1_*X*_+2_*X*_+3_ and 500 members of C-terminal *X*_−3_*X*_−2_*X*_−1_-*p*S-*P*_+1_*P*_+2_*P*_+3_ sublibraries each. Peptide residues in the phosphopeptide fragments were designated as P and X, then numbered at subscript according to their proximity to the phosphoserine and negative if they are located N-terminal of it. Ten representative amino acids (Ala, Glu, Phe, Gly, Lys, Leu, Pro, Gln, Arg, Val) were used at the position of *P*±_1/2_, and among them five amino acids (Glu, Phe, Leu, Pro, Arg) at position of *P*±_3_. X represents an isokinetic mixture of rest 14 amino acids. The relative fluorescence values were generated by fluorescence signal of Cy3-labeled 14-3-3s, which represented the affinity between phosphopeptide and 14-3-3s. To improve the data discrimination, raw relative fluorescence values were transformed by taking the logarithm to the base of two before analysis [58]. The processed data represented the relative binding affinity of 14-3-3 isoforms with each phosphopeptide fragment.

### 4.2 Amino acids descriptors

The tripeptide sequences in the phosphopeptide fragments were extracted and used for the rest of the analysis. The relative binding affinity of each tripeptide can be characterized by describing the residues with descriptors. Nonbonding effects, like electrostatic, van der Waals, hydrophobic interactions and hydrogen bond play central roles in phosphopeptide interaction with all type of 14-3-3 isoforms [43]. DPPS descriptor is a kind of suitable way to be used as the descriptor here [48]. The DPPS descriptors were made up of ten variables for describing the electronic property, steric property, hydrophobicity and hydrogen bond of all amino acids (Tab.2). For tripeptide sequences in both N- and C-terminal sublibraries, physicochemical parameters of amino acid residue at each position were characterized by 10 DPPS descriptors. By assigning DPPS to tripeptide, totally of 30 variables were generated. Due to the different contributions of residue position and its physicochemical parameters to binding affinity, variable selection is required prior to QSAR modeling, expecting to improve modeling qualities and to decrease complexity. The key variables for the binding affinity were screened by stepwise regression. Then based on the screened variable, we can build a QSAR model for each sub-library of seven isoforms.

**Table 2:** Amino acids descriptors

| AA | V1 | V2 | V3 | V4 | V5 | V6 | V7 | V8 | V9 | V10 |
| --- | --- | --- | --- | --- | --- | --- | --- | --- | --- | --- |
| Ala | -1.02 | -2.88 | -0.56 | 0.36 | -6.15 | -1.68 | 0.04 | -2.51 | -1.94 | -0.01 |
| Arg | 1.99 | 4.13 | -4.41 | -1.02 | 4.78 | 3.04 | -9.06 | 6.71 | 4.41 | 0.07 |
| Asn | -2.19 | 1.86 | 0.38 | -0.13 | -2.30 | 1.41 | -5.71 | -1.11 | 1.73 | -0.19 |
| Asp | -6.60 | 3.32 | 1.61 | 0.36 | -3.25 | 1.95 | -7.36 | 0.14 | 1.24 | -0.15 |
| Cys | 0.21 | 1.12 | 3.42 | -0.68 | -2.27 | -1.22 | 3.11 | -2.98 | -1.70 | 1.57 |
| Gln | -0.47 | 1.16 | -0.57 | 0.69 | 0.39 | 1.93 | -5.46 | -0.84 | 1.93 | 0.85 |
| Glu | -5.39 | 0.65 | -0.98 | 1.39 | -0.23 | 2.51 | -6.84 | -0.68 | 1.41 | 1.28 |
| Gly | -2.86 | -5.00 | -2.97 | 0.53 | -11.45 | 1.89 | -2.11 | -3.99 | -2.16 | -0.76 |
| His | 0.73 | 2.68 | -0.66 | -1.89 | 1.60 | 1.13 | -1.94 | -0.11 | 0.44 | 0.15 |
| Ile | 1.91 | -3.13 | 0.01 | 1.14 | 2.70 | -4.55 | 8.93 | 0.18 | -1.10 | -0.76 |
| Leu | 1.64 | -2.57 | 0.00 | 1.35 | 2.62 | -2.65 | 7.72 | 0.05 | -1.03 | -1.81 |
| Lys | 2.47 | 1.54 | -4.28 | -0.86 | 2.77 | 2.06 | -6.18 | 2.05 | 2.19 | -1.65 |
| Met | 1.93 | -0.01 | 1.21 | 0.99 | 2.79 | -0.56 | 5.33 | -0.87 | -0.99 | -1.09 |
| Phe | 2.68 | 0.84 | 2.22 | 0.71 | 5.02 | -0.30 | 8.60 | 1.13 | -1.40 | -0.28 |
| Pro | 0.45 | -2.89 | 1.77 | -5.81 | -3.79 | -0.61 | 0.70 | 1.21 | -1.67 | 1.79 |
| Ser | -1.76 | -0.19 | 1.06 | -0.69 | -5.72 | 0.14 | -4.14 | -2.42 | -0.13 | 0.69 |
| Thr | -0.55 | -0.66 | 0.13 | -0.31 | -2.76 | -1.56 | -2.46 | -2.12 | 0.17 | 0.08 |
| Trp | 3.88 | 1.78 | 1.68 | 2.00 | 9.31 | 0.89 | 7.53 | 4.27 | -0.23 | -1.42 |
| Tyr | 2.10 | 1.26 | 1.15 | 0.91 | 5.90 | 0.74 | 3.71 | 3.32 | 0.25 | 1.33 |
| Val | 0.83 | -3.02 | -0.22 | 0.97 | 0.05 | -4.55 | 5.61 | -1.41 | -1.44 | 0.30 |

Electronic property: columns V1-V4, steric property: columns V5-V6, hydrophobicity: columns V7-V8, hydrogen bond: columns V9-V10.

### 4.3 QSAR modelling

The initial dataset is divided into two parts, one for training and the other for testing. For example, there are 500 relative affinity data in one of isoform sublibraries. 400 samples of the relative affinity data are randomly selected as training dataset and the rest of 100 samples as testing dataset. The training datasets are used for model construction and testing datasets are used for performance test. After construction of modeling finished, the test datasets are used for the confirmations of the performance. Linear regression is the commonly used modeling methods for QSAR studies. It has advantages in its simplicity, accuracy and ease of interpretation. Hence linear regression is chosen for modeling here.

The model assumes that there is a linear relationship between the peptide activity and the variables we extracted by the amino acid descriptor. For each peptide sequence, the relative binding affinity value *f* (*X*) is represented by a vector of *p* variables:

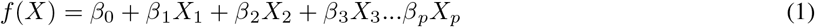

where *β*_0_ is the model constant, *X*_1_, *X*_2_, *X*_3_…*X_P_* are a series of amino acid descriptors with their corresponding coefficients *β*_1_, *β*_2_, *β*_3_…*β_p_* based on the regression setting. Once the tripeptide sequence was translated into a two-way data matrix by amino acid descriptors, regression methods such as ordinary least squares can be used to develop the QSAR models relating sequence to bioactivity.

The panelized regression methods, including lasso, ridge and elastic net regression, can create the linear models that is panelized, that is, a constraint *λ* is added in the equation to minimize the sum of squared residuals[5]. The consequence of importing this penalty, is to reduce the coefficient values towards zero. This allows variables with less contributions to have a coefficient close to or equal zero. The main difference between lasso and ridge regression is the penalty term they use. Ridge uses *L*_2_-norm which is the sum of squared coefficients, 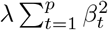. Lasso uses *L*_1_-norm which is the sum of the absolute coefficients, 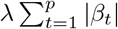. *L*_1_-norm imposes sparsity among the coefficients, which makes the fitted model more interpretable [15]. Then elastic net was introduced and has a penalty which is a mix of *L*_1_- and *L*_2_-norms, 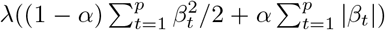. Since *α* is the mixing parameter between ridge(*α* = 0) and lasso (*α* = 1), elastic net is regarded as a compromise between lasso and ridge. Here, elastic net regression was selected to establish the QSAR models since it solves the limitations of both lasso and ridge. The 10-fold cross-validation methods were used to choose the tuning parameter *λ* and *α*, which is able to minimize the cross validation error. The training dataset is divided randomly into ten parts and then each of parts are used as testing for the model on the other parts. Then the best value pair for *λ* and *α* was selected for the resulting models.

To get the measures of goodness-of-fit for QSAR models, the Pearson’s product-moment correlation coefficient squared (*R*^2^) and the root mean squared error (*RMSE*) are calculated between prediction and real dataset. RMSE can be calculated by

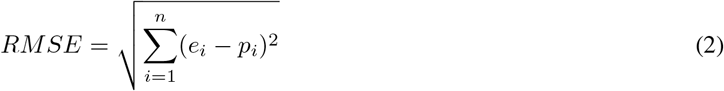

where *e_i_* is *i*-th binding affinity value from the experiment dataset with *n* samples, *p_i_* is *i*-th value from the predicted results by the models. And *R*^2^ can calculated by

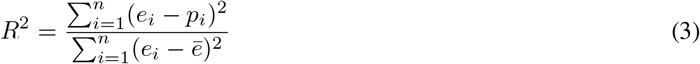

where 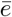 is the average binding affinity from the experimental dataset. Ideally, lower *RMSE* and higher *R*^2^ values are indicative of a good model. The most ideal result would be an *RMSE* value of zero and *R*^2^ value of 1.

### 4.4 Data analysis

Cluster analysis of the relative binding affinity was performed using euclidean measure to obtain distance matrix and complete agglomeration method for clustering (*heatmap.2* in R package *gplots*). The relative binding affinity values were scaled before the analysis. The sequence conservation analysis was performed by uploading the peptide sequences to weblogo web server [7]. The default parameters were used in this analysis.

## 5 Conclusions

In this study, the development of the predictive QSAR models and prediction of the binding affinity of phosphopeptide fragments against 14-3-3s were described. According to the models, the important properties of residues in the phosphopeptide were identified. The conserved patterns of peptide sequences were verified in our predicted results. The well-known ligands against 14-3-3s were also found in our study. The predictions provide the evidences that the conserved peptides existed in the 14-3-3s interaction and could be applied in the further studies.

14-3-3s are regarded as the novel therapeutic targets for cancer therapy. A small portion of positive charged residues located in the characterized phosphopeptide binding domain in 14-3-3s. This domain is required for binding and makes 14-3-3s an ideal molecular target. Currently, there are not a small molecule inhibitor of 14-3-3s developed in the clinic. It will be important to elucidate the underlying mechanism of 14-3-3s interaction that could be mined in cancer study by the efforts from computational and biological works.

